# A COVID-19 Rapid Antigen Test Employing Upconversion Nanoparticles

**DOI:** 10.1101/2024.08.05.606725

**Authors:** Le Zhang, Jiajia Zhou, Olga Shimoni, Shihui Wen, Amani Alghalayini, Yuan Liu, Meysam Rezaeishahmirzadi, Jiayan Liao, Mahnaz Maddahfar, Roger Hunt, Murdo Black, Matt D. Johansen, Phil M. Hansbro, Lin Zhang, Martina Stenzel, Majid Warkiani, Stella M Valenzuela, Dayong Jin

**Author notes:** These authors contributed equally to this work (Le Zhang, Jiajia Zhou).

## Abstract

The COVID-19 pandemic has underscored the critical need for rapid and accurate diagnostic tools. Current methods, including PCR and rapid antigen tests (RAT), have limitations in speed, sensitivity, and the requirement for specialized equipment and trained personnel. Nanotechnology, particularly upconversion nanoparticles (UCNPs), offer a promising alternative due to their unique optical properties. UCNPs can convert low-energy near-infrared (NIR) light into higher-energy visible light, making them ideal for use as optical probes in single molecule detection and point of care applications. This study, initiated in early 2020, explores the opportunity of using highly doped UCNPs (40%Yb^3+^/4%Er^3+^) in lateral flow assay (LFA) for the early diagnosis of COVID-19. The UCNPs-based LFA testing demonstrated a minimum detection concentration of 100 pg/mL for SARS-CoV-2 antigen and 10^5^ CCID_50_/mL for inactivated virus. Clinical trials, conducted in Malaysia and Western Australia independently, showed that the technique was at least 100 times more sensitive than commercial RAT kits, with a sensitivity of 100% and specificity of 91.34%. The development process involved multidisciplinary collaborations, resulting in the Virulizer device, an automated strip reader for point-of-care testing. This work sets a reference for future development of highly sensitive and quantitative rapid antigen tests, aiming for the Limits of Detection (LoD) in the range of sub-ng/mL.

## Introduction

At the onset of the COVID-19 pandemic, early detection was crucial due to the rapid transmission of the virus. However, PCR testing has a relatively slow turnaround time, and the standard rapid antigen test (RAT) using gold colloidal nanoparticles were not sensitive, which hinders the timely control of the outbreak spread [1]. While nasopharyngeal swabs, as the standard sampling method, cause discomfort and risk to patients [2], saliva sampling is more convenient and non-invasive [2] with higher acceptance and operability, allowing patients to collect their saliva samples themselves, thus reducing dependency on medical resources and lowering the risk of cross-infection [2]. However, the viral concentration in saliva (10^4^-10^6^ copies/mL which equals to 30 fg/mL to 3 ng/mL of nucleoprotein lysed in the saliva as viral antigen) is generally lower than in nasopharyngeal swab samples, posing a significant challenge to the accuracy of conventional rapid test strip technology [3].

UCNPs can convert low-energy near-infrared (NIR) light into higher-energy visible emissions [4]. This property offers minimal background interference [4,5]. Typically, UCNPs are β-NaYF_4_ nanoparticles co-doped with 20 mol% Yb^3+^ and 2 mol% Er^3+^ [6]. To increase the brightness of UCNPs, we developed a series of highly doped UCNPs that exhibit a nonlinear enhancement in luminescence intensities, depending on the excitation power density [7]. Before the pandemic, we successfully applied the highly doped UCNPs in a lateral flow assay (LFA) and achieved quantitative measurement of cancer biomarkers [8].

To address the emergency and global demand for highly sensitive and rapid detection methods, in early 2020, we initiated a transformational project with the industry partner Alcolizer, an Australian company who has extensive market development experience in saliva testing for illicit drugs, aiming to translate UCNPs-based lateral flow assays into a more sensitive COVID-19 RAT. The project was focused on the saliva test, including saliva collection, viral lysis, antigen enrichment and LFA using highly doped UCNPs. While achieving high detection sensitivity, towards a point of care test (POCT), we also made significant progress in the development of a detection device using a laser diode and a CCD detector, initially from a lab-based optical microscope, followed by a series of iterations and industry designs into a small-footprint automated strip reader under the trade name of Virulizer.

The validation of this study has undergone three stages, including the laboratory proof-of-concept, preclinical validation using inactivated SARS-CoV-2 virus, and final independent clinical trials. Initially, laboratory validation using purified SARS-CoV-2 antigen resulted in a minimum detection concentration close to 100 pg/mL. By mid 2021, collaborating with the Centenary Institute, Centre for Inflammation at UTS and The Prince of Wales Hospital in New South Wales, Australia, we conducted two independent validation studies using inactivated SARS-CoV-2 virus in a PC3 laboratory environment. Both results confirmed the feasibility of our method, with a minimum detectable virus concentration close to 10^5^ CCID_50_/mL. By early 2022, a first-round clinical trial in Malaysia was conducted [9]. The results showed that our method was at least 100 times more sensitive than the commercially available RAT kits on the market. Later, in June 2022, an independent research team conducted the second round of clinical trials at the PathWest COVID-19 Drive-Through Clinic (DTC) in Western Australia. The results further demonstrated the high sensitivity and specificity of UCNPs-based RAT for diagnosing early-stage COVID-19 patients, with sensitivity reaching 100% and specificity reaching 91.34% [not yet published].

The process of developing the UCNPs-based COVID-19 RAT involves a set of multidisciplinary research collaborations and has attracted considerable media attention [10,11] and public awareness [12]. Here, we provide the details of this research project, along the progress of each critical milestones, including our experimental design and our key findings. Our work sets a reference for future development of quantitative and ultrasensitive RATs for point of care and point of risk tests, that requires the limit of detections (LoDs) in the range of below ng/mL.

## Materials and Methods

### Materials

Yttrium(III) chloride hexahydrate (YCl_3_·6H_2_O), ytterbium(III) chloride hexahydrate (YbCl_3_·6H_2_O), erbium(III) chloride hexahydrate (ErCl_3_·6H_2_O), sodium hydroxide (NaOH), ammonium fluoride (NH_4_F) with 99.99% trace metals basis, oleic acid (OA), 1-Octadecene (ODE) with technical grade 90%, ethanol, methanol 100% pure, cyclohexane for HPLC, ≥99.9%, tetrahydrofuran (THF) anhydrous, ≥99.9% inhibitor-free, monomer poly(ethylene glycol) methyl ether methacrylate (EGMEMA) with average Mn 500 g/mol, contains 100 ppm MEHQ as inhibitor, 200 ppm BHT as inhibitor, 4-Cyano-4-(phenylcarbonothioylthio)pentanoic acid (CPADB, RAFT agent), 2,2’-azobisisobutyronitrile (AIBN) 97%, purified by recrystallization twice from methanol, monomer ethylene glycol methacrylate phosphate (EGMP), n-hexane for HPLC ≥99.9%, toluene, Milli-Q integral water, dialysis membrane with various molecular weight cut-offs with 3.5 kDa were purchased from Sigma Aldrich. Particle Conjugation Kit was supplied from AnteoTech company, Australia. Anti-nucleoprotein monoclonal antibody (NUN-S91) was purchased from Sino Biological.

### Synthesis of highly doped UCNPs

NaYF_4_: Yb^3+^, Er^3+^ nanocrystals (core) with high doping (4%Er/40%Yb) were synthesized according to our previously reported method. Briefly, 1 mmol RECl_3_·6H_2_O (RE = Y, Yb, Er) with the mole ratio of (Y = 56%, Yb = 40 %, Er = 4%) was mixed together in a 50 mL-three necked round bottom flask containing 6 mL OA and 15 mL ODE. The mixture was heated to 160 °C under argon flow for 30 min to obtain a clear solution and then cooled down to room temperature, and 5 mL methanol solution of NaOH (2.5 mmol) and NH_4_F (4 mmol) was added, followed by 30 min stirring at room temperature. The reaction mixture was heated to 120 °C under argon flow for another 20 min to remove the methanol, then the solution was further heated to 300 °C for 90 min. Finally, the reaction solution was cooled down to room temperature, the product was precipitated by ethanol followed by centrifugation at 7600 g for 6 min. The final UCNPs were washed 3 times by cyclohexane and OA as well as ethanol and methanol followed by centrifugation at 7600 g for 6 min.

To get the nanoparticles with core-shell structure, layer-by-layer epitaxial growth has been employed. The shell precursor’s preparation is similar with that for the core nanoparticles synthesis, until the step where the reaction solution was slowly heated to 150 °C and kept for 20 min. Instead of further heating to 300 °C to trigger nanocrystal growth, the solution was cooled down to room temperature to yield the shell precursors (NaYF_4_). For epitaxial growth, 0.15 mmol as-prepared core nanocrystals were added to a flask containing 6 mL OA and 6 mL ODE. The reaction mixture was heated to 170 °C under argon flow for 30 min, and then further heated to 300 °C. Next, 0.25 mL as prepared shell precursors were injected into the reaction mixture and ripened at 300 °C for 4 min, followed by the same injection and ripening cycles for 3 min interval to get the nanocrystals with the desired size. Finally, the slurry was cooled down to room temperature and the formed nanocrystals were purified according to the same procedure used for the core nanocrystals.

### Synthesis of PEGMEMA-*b*-PEGMP diblock copolymer

The RAFT agent used in our experiment was 4-Cyano-4-(phenylcarbonothioylthio)pentanoic acid (CPADB). Synthesis of PEGMEMA-*b*-PEGMP diblock copolymer was employed through Reversible Addition Fragmentation Chain Transfer (RAFT) polymerization. In brief, EGMEMA (8g, 2.67×10^−2^ mol), CPADB (36.4 mg, 1.3×10^−4^ mol), and AIBN (2.13mg, 1.3×10^−5^ mol) were dissolved in toluene (9 mL), the reaction mixture was placed in a round-bottom flask equipped with a magnetic stirrer bar. The flask was then sealed with a rubber septum and purged with nitrogen gas for 60 min. The reaction mixture was then placed in a preheated oil bath at 70 °C. To prepare PEGMEMA macro-RAFT with 15 monomer repeats, the polymerization was terminated after 3.5 h by placing the reaction solution into an ice bath for 15 min. The synthesised PEGMEMA macro-RAFT was purified by adding excess n-hexane followed by centrifugation at 7600 g for 5 min, then the polymer was dissolved in the methanol and then was dialysed (3.5 KDa cut-off) against 500 mL methanol for 48 h. The final sample was dried under vacuum oven at 40 °C for 14 h followed by storing at 4 °C until required for further chain extension to form diblock copolymer. PEGMEMA was used as a macro-RAFT agent for chain extension with EGMP to introduce the functional phosphonic acid group that allows coordination with lanthanide ions on the surface of UCNPs. PEGMEMA (5 × 10^−5^ mol), EGMP (3 × 10^−4^ mol) and AIBN (5×10^−6^ mol) were dissolved in methanol (1 mL) in a round bottom flask equipped with a magnetic stirrer bar. The reaction mixture was degassed with N_2_ gas for 45 min in an ice bath. The polymerization was carried out in a preheated oil bath for 17 h at 70 °C. The reaction was terminated by placing the sample in an ice bath for 15 min when the monomer conversion is around 90%. The monomer conversion measurement was conducted with H NMR [13]. The diblock copolymer was purified by dialysis (3.5 kDa cut-off) against methanol for 4 h two times to remove unreacted EGMP, and the purified polymer was then dried in a vacuum oven for 14 h to remove the remaining solvent. The presence of phosphoric group of EGMP (second block) was confirmed by ^1^H NMR and ^31^P NMR.

### Surface modification of UCNPs with PEGMEMA-*b*-PEGMP

500 *μ*L of OA-coated UCNPs (20 mg/mL) kept in cyclohexane were centrifuged at 20,240 g for 30 min and redispersed in 1 mL THF. A solution of the diblock copolymer (10 mg) in 500 *μ*L THF was individually added to UCNPs dispersion. The reaction solution was sonicated for 1 min followed by incubation in a shaker for 14 h at room temperature. The polymer-coated UCNPs were purified four times by washing with different volume/ratio of THF/MiliQ water followed by centrifugation at 20,240 g for 30 min. Finally, the polymer-coated UCNPs were then resuspended in 1 mL MiliQ water for further use.

### Bioconjugation of polymer coated UCNPs with anti-nucleoprotein monoclonal antibody (UCNPs@mAb)

The bioconjugation of UCNPs with antibodies using the Particle Conjugation Kit (AnteoBind-Nano Kit for particles < 0.5 *μ*m) proceeded according to the manufacturer’s protocol (AnteoTech, Australia). The kit contains particle activation solution, particle wash solution, conjugation buffer at pH= 6, blocker diluent at pH= 6, and storage buffer at pH= 8. In brief, 50 *μ*L polymer-coated UCNPs (20 mg/mL) were mixed with the particle activation solution (450 *μ*L) followed by shaking at 700 rpm at room temperature for 1 h. 200 *μ*L particle washing solution was added to the reaction followed by centrifugation at 20,240 g for 1 h. The nanoparticles were redispersed in 200 *μ*L conjugation buffer followed by centrifugation at 20,240 g for 1 h. The activated polymer-coated nanoparticles were resuspended in 200 *μ*L conjugation buffer, then, 250 *μ*L anti-nucleoprotein monoclonal antibody (NUN-S91, 50 *μ*g) at concentration of 0.2 mg/mL was added to the activated nanoparticle’s solution followed by incubation at room temperature for 1 h. After that, 50 *μ*L blocking buffer was added to UCNPs@mAb solution followed by 1 h incubation at room temperature to block unreacted ligands. The solution was centrifuged at 20,240 g for 1 h and redispersed in 200 *μ*L storage buffer. This washing step was repeated twice and final UCNPs@mAb were resuspended in 100 *μ*L storage buffer and stored at 4 °C for further use.

### Characterization of UCNPs

The morphology characterization of the nanoparticles was performed by transmission electron microscope (TEM). The FEI Tecnai T20 microscope is operated under an accelerating voltage of 200 kV. The cyclohexane dispersed UCNPs were imaged by dropping them onto carbon-coated copper grids.

### Fabrication of lateral flow assay strips

The paper-based strip consists of an adhesive PVC back pad, a nitrocellulose membrane (FF120HP membrane, GE Life Science), a sample pad (CF4, GE Life Science) and an absorbent pad (CF5, GE Life Science). The test area was printed by 0.5 *μ*L of 0.2 mg/mL anti-nucleoprotein polyclonal antibodies produced in mice (NUN-S92, Sino Biological) using XYZ platform dispenser (HM 3260, Shanghai Kinbio Tech). The NC membrane was then dried in a desiccator overnight (14 h) at room temperature. The sample pad, conjugation pad, nitrocellulose membrane and absorbent pad were orderly mounted on the PVC back pad with a 2 mm overlap between each two adjacent pads. The assembled pads were cut into strips with a width of 3 mm with programmable shear (Samkoon, Gold bio).

### Lateral flow assay for detecting antigen molecules in saliva

We used a medical-grade sponge to collect saliva. Saliva samples were diluted in a homemade RIPA lysis buffer composed of 100 mM Tris (pH= 7.6), 300 mM NaCl, 0.1% SDS, and 1% NP-40. The mixture was gently agitated to ensure thorough mixing and then incubated for 5 minutes at room temperature. Then, 50 *μ*L of the diluted sample was loaded into the sample port of an LFA strip. As the liquid migrated from the sample pad to the absorbent pad, the bioconjugate of functionalized UCNPs@mAb were captured at the test line. After a 10-minute incubation period, the strip was inserted into the strip reader to detect the UCNPs signal at emission wavelength at 653 nm with the signal peak in the test line.

### Data and statistical analysis

Data were analysed using two-tailed unpaired T-tests or one-way ANOVA with post-hoc multiple comparison tests in GraphPad Prism 10.0. A *p*-value of less than 0.05 was considered statistically significant.

### Results and Discussion

The development of an upconversion immunoassay for the detection of SARS-CoV-2 viral antigens critically depends on the design of the luminescent reporter that replaces the conventional enzyme amplification steps. Here, we use the precipitation method to yield high quality monodispersed core-shell UCNPs, which is governed by a temporal nucleation and growth of nanocrystals. The TEM characterization results in Figs. 1A and 1B show one batch of core-shell UCNPs used during the development of our technology. The TEM image in Fig. 1A reveal the uniformity in morphology, as well as the monodispersity of UCNPs in cyclohexane. UCNPs were coated with a copolymer that possesses a terminal carboxylic acid functional groups due to the RAFT agent. The carboxyl groups improve particle dispersion in water as well as serving as attachment sites for subsequent antibody conjugation steps [14]. The copolymer layer is difficult to observe in the TEM characterization results shown in Fig. 1B due to the low contrast of the involved elements. However, the monodispersity of the modified UCNPs in water is apparent from the TEM images. We optimized the designs of copolymers with varying repeats of EGMEMA (Table 1), as well as the modification protocols for each type of copolymer to modify the UCNPs. We found that their surface charges vary significantly depending on the number of EGMEMA repeats (Table 1). Shorter polymers result in denser polymer brushes, thus higher COOH density [14]. We concluded that copolymers with 15 or 17 repeats of EGMEMA are preferred for this work, as they provide the highest zeta potential (surface charge), indicating the highest degree of stability for the UCNPs (Figure 2). They also offer sufficient -COOH groups for subsequent bioconjugation (Fig. 2).

**Table 1.** Zeta potential of different diblockcopolymer coated UCNPs.

| Polymer type ( different repeats of EGMEMA units) | Zeta Potential (mV) |
| --- | --- |
| 15 repeats | From -20 to -28 |
| 17 repeats | From -22 to -31 |
| 26 repeats | From -10 to -12 |
| 60 repeats | From -6 to -15 |
| 80 repeats | From -4 to -6 |

**Figure 1.**
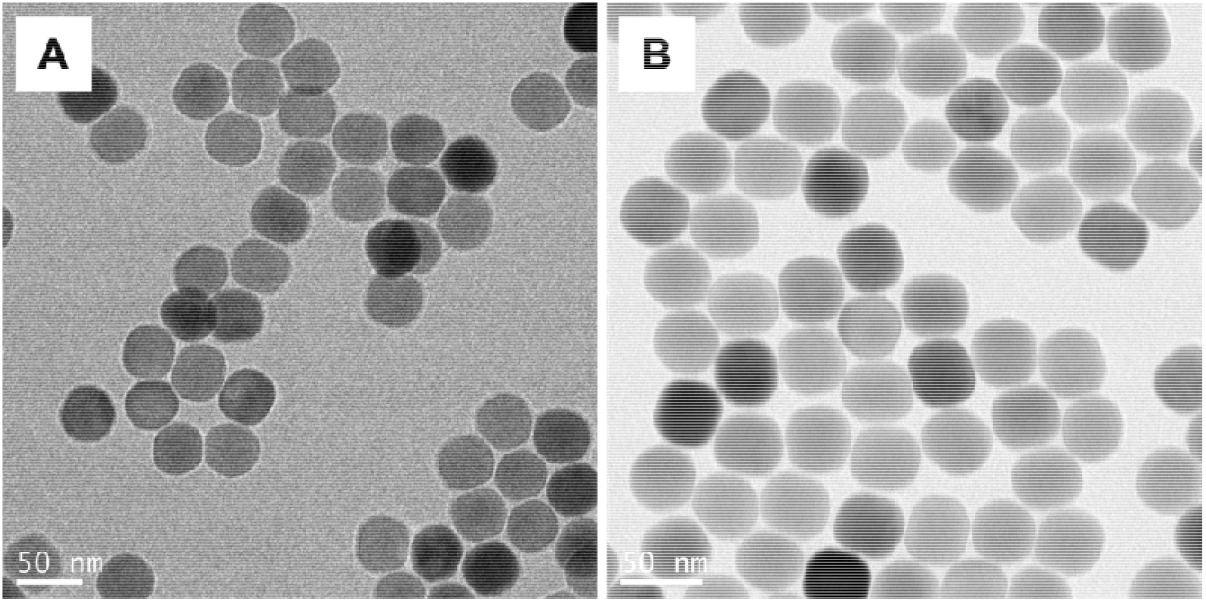
Uniform morphology and surface modification of highly doped UCNPs. **(A)** TEM images of NaYF_4_: 40%Yb,4%Er@NaYF_4_ before the surface modification. **(B)** TEM images of NaYF_4_: 40%Yb,4%Er@NaYF_4_ after the surface modification of PEGMEMA_60_-*b*-PEGMP_3_ copolymer.

**Figure 2.**
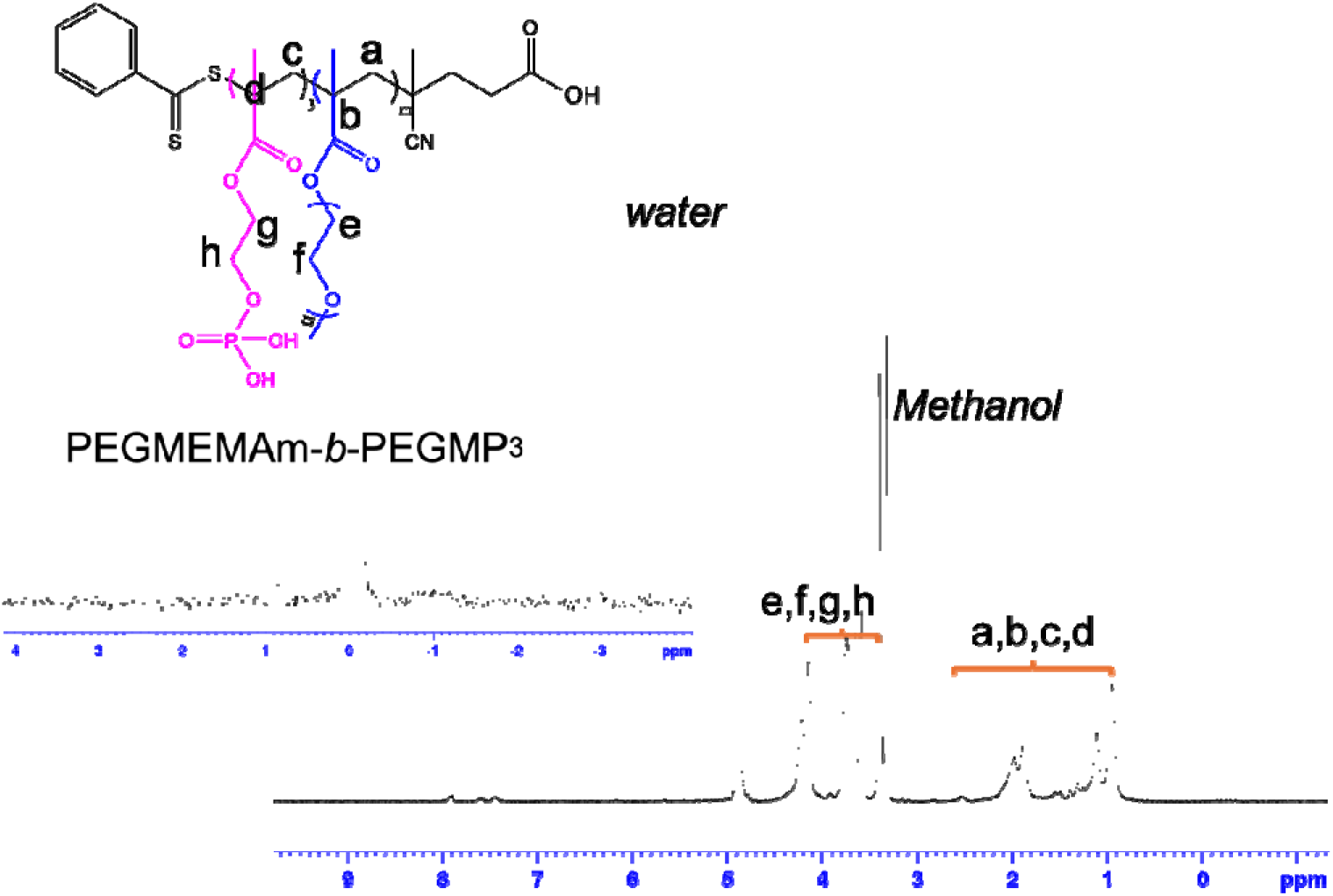
PEGMEMA_15_-*b*-PEGMP_3_ diblock-copolymer. ^1^H NMR and ^31^P NMR spectra of pure PEGMEMA_15_-*b*-PEGMP_3_ in Methanol-d_4_ (MeOD).

After the polymer modification, the polymer-coated UCNPs were then conjugated to anti-nucleoprotein monoclonal antibody by using a commercial kit (AnteoTech, Australia). Traditional LFA test strips rely on a control line to confirm the proper flow of reporter molecules, which must be detected in a highly concentrated or aggregated state, such as with gold or europium nanoparticles. Here, UCNPs emit luminescence strong enough to be detected without the need for aggregation or control line [15,16]. As a result, for UCNPs, the flow of the test strip can be verified by detecting any position under high density laser excitation.

To read the luminescence signal of UCNPs on the strip, a series of detection systems with a 980 nm continuous-wave laser diode have been designed and validated. In early 2020, we initially designed the first-generation imaging device following a typical design of epi-fluorescence optical layout, with the size of footprint getting smaller and smaller (three aluminium mechanically engineered units and one 3D printed plastic unit shown in Fig 3A). By the end of 2020, we developed a table-top unit with improved mechanical stability and optical safety (Fig. 3B). The unit employs an Ocean Optics USB2000 spectrometer as the detector and a 980 nm fibre coupled laser diode as the excitation source. In 2021, a design of a hand-held unit has been achieved with a computer pad still needed (Fig 3C). The final design of a Virulizer reader with a CMOS camera chip used for signal detection, a user-friendly touchscreen, and an internet unit for data communication, has been achieved, as displayed in Fig. 3D. For the optics design, briefly, the incident 980 nm laser first passed through a 920 nm long-pass filter (Filter #1) to eliminate unwanted emissions in visible range from the diode laser. Subsequently, a lens was employed to collimate the beam before being reflected by a dichroic mirror and focused by an objective lens into a focal spot with a diameter of 0.4 mm on the strip testing area. The emitted upconversion luminescence from the highly doped UCNPs at the two peaks of around 528-545 nm and around 640-660 nm was collected by the objective lens, passing through the dichroic mirror and a dual-bandpass filter with 524 nm and 673 nm center wavelengths (Filter #2), being focused and captured by a CMOS camera (Fig. 3A).

**Figure 3.**
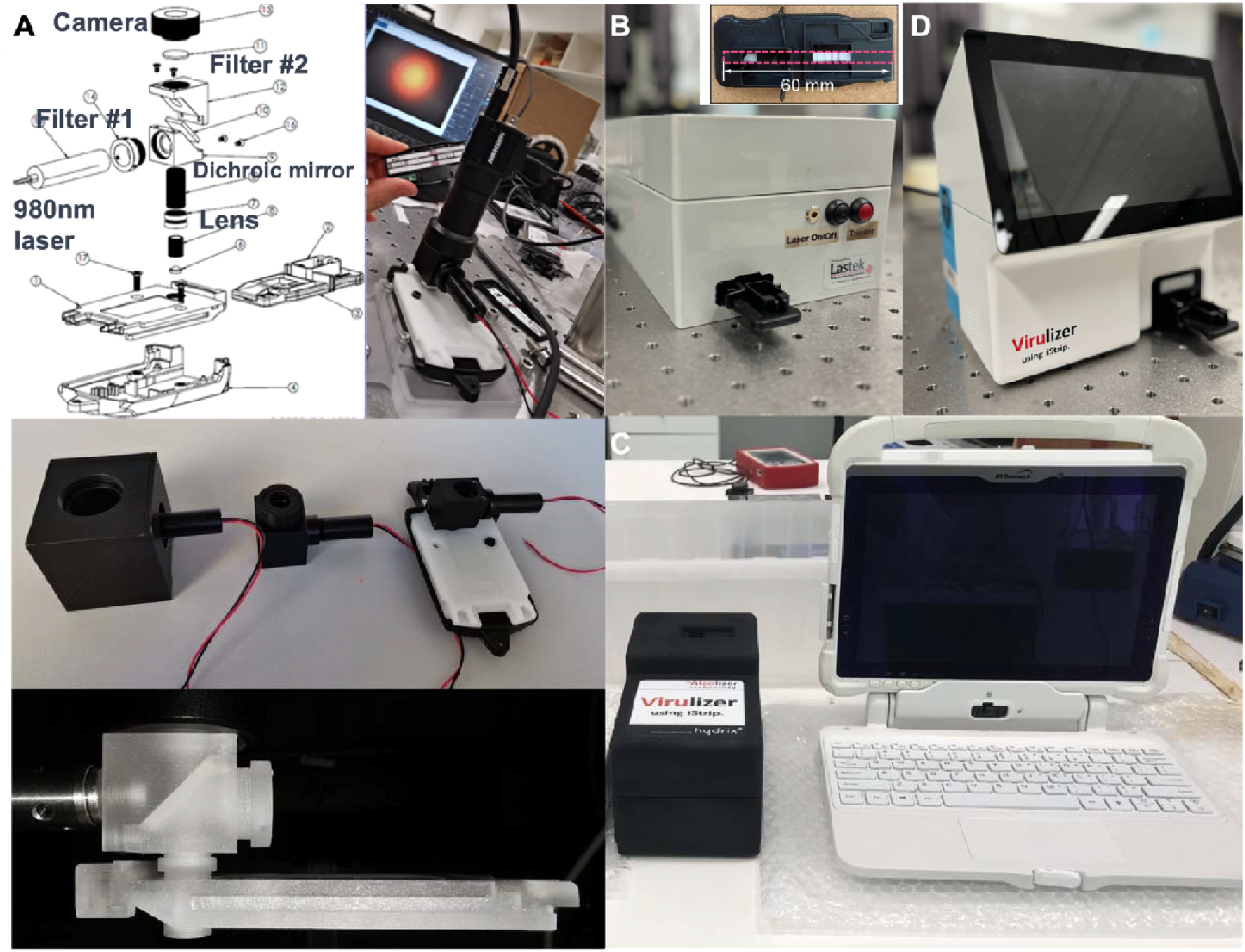
Four generations of test strip readers have been developed. This includes the first generation of optical excitation and signal collection devices with three representative sizes of mechanically engineered aluminium units and one 3D printed plastic unit to match the strip holder **(A)**, the table-top unit **(B)**, a hand-held unit with a computer pad **(C)**, and the final Virulizer reader **(D)**.

To optimize the reading time and the signal along the strip position, we first mixed inactivated SARS-CoV-2 virus (virus loads at 1×10^5^ CCID_50_/mL) with RIPA buffer (*v/v*, 1:1), and then applied 50 *μ*L of the mix to the test strip. By scanning the distribution of luminescent signals at different positions on the strip at various time points, as shown in Fig. 4, the highest signal was identified at the point of 15 minutes after the sample loading with a notably significant specific positive signal for antigen detection achieved.

**Figure 4.**
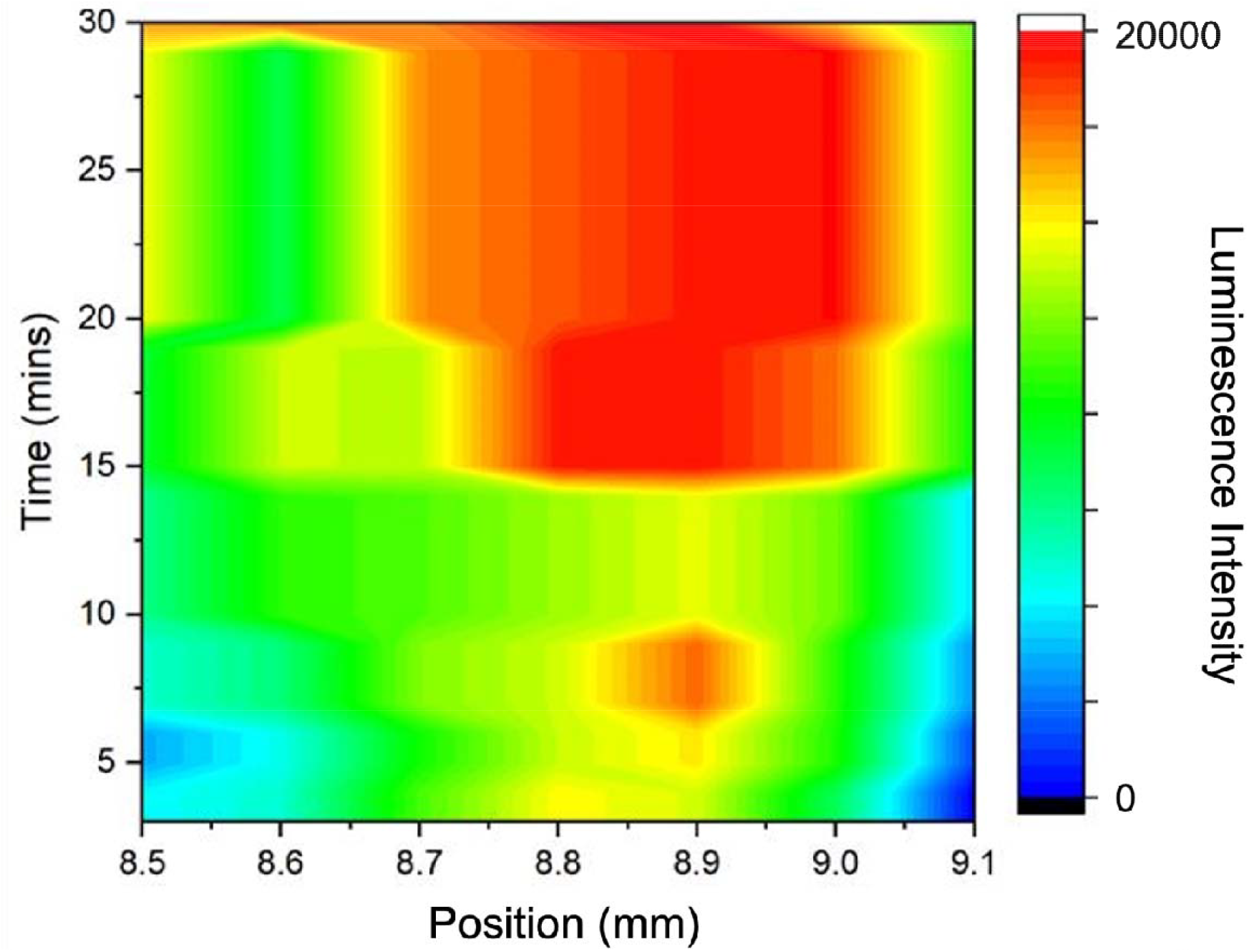
The heat map of luminance intensities around the test line (over a width of 0.6 mm) and time points after the antigen sample loading (3 to 30 min).

The performance of the UCNPs-based LFA was evaluated using the purified SARS-CoV-2 nucleoprotein antigen and inactivated SARS-CoV-2 virus, respectively. A serial dilution of the purified SARS-CoV-2 nucleoprotein in RIPA lysis buffer was prepared at concentrations of 0, 1, 10, 10^2^, 10^3^, 10^4^, 10^5^, 10^6^, and 10^7^ pg/ml and 50 *μ*l volume of sample was applied onto the sample pad on each strip. The lowest detectable concentration of the nucleoprotein was found around 10^2^ pg/mL (Fig. 5A). Furthermore, the inactivated SARS-CoV-2 virus was lysed by RIPA buffer to extract the nucleoprotein antigen and diluted in RIPA lysis buffer (*v/v*, 1:1) to prepare dilutions of 0, 3.3 × 10^4^, 2.1 × 10^5^, 4.1 × 10^5^, 1.25 × 10^6^, 1.7 × 10^6^, 2.5 × 10^6^, 3.3 × 10^6^ CCID_50_/ml (viral concentrations), and the lowest detectable concentration of the inactivated virus was found around 3.3 × 10^4^ ∼ 2.5 × 10^5^ CCID_50_/ml (Fig. 5B).

**Figure 5.**
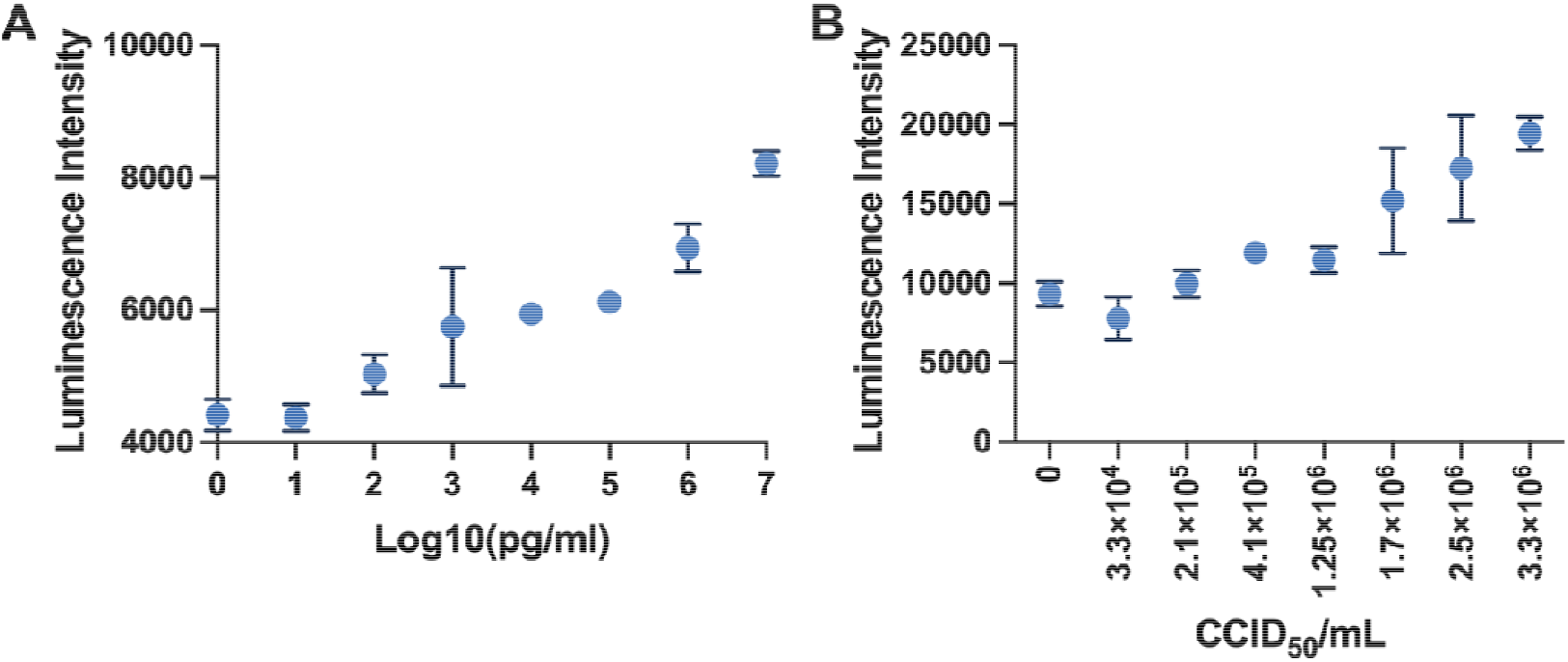
Performance of UCNPs-based LFA. **(A)** at different concentrations of SARS-CoV-2 nucleoprotein in RIPA buffer. **(B)** at different concentrations of inactivated SARS-CoV-2 virus lysed in RIPA buffer. Each experiment was independently repeated six times using parallel replicates, and the data are shown as mean ± SEM.

Finally, we tested the performance of UCNPs-based LFA strip by using saliva containing inactivated SARS-CoV-2 virus. This method replicates the viscosity and composition of saliva and mucus produced by the salivary glands in the mouth. The saliva was spiked with inactivated SARS-CoV-2 virus to simulate varying viral loads from patient samples. It was shown that this assay had a high sensitivity in the detection of inactivated virus in saliva containing inactivated SARS-CoV-2 virus at 1 × 10^5^ CCID_50_/ml, which approximates the viral load typically present in the saliva of COVID-19 patients during early infection stages (Fig. 6) [3,17].

**Figure 6.**
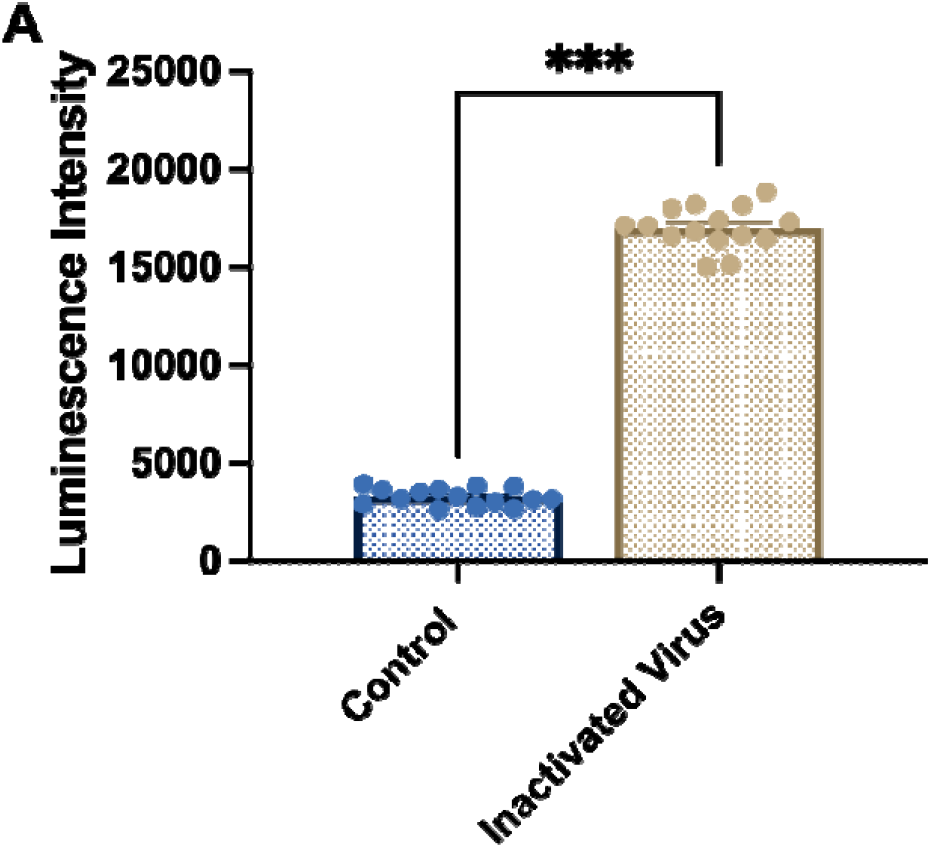
Performance for the detection of inactivated SARS-CoV-2 virus mixed with saliva sample. Luminescence intensities of testing areas for the detection of control and inactivated SARS-CoV-2 virus mixed with saliva samples. Each experiment was independently repeated at least 15 times using parallel replicates, and the data are shown as mean ± SEM. Significant differences among different groups are analysed by independent *t*-test, \*\*\**P* < 0.001.

The advancements presented in this study hold significant implications for public health. The development of more efficient and accurate RAT methods is crucial for controlling the spread of the virus, leading to more effective quarantine measures (Fig. 7). The much lower LoD guarantees a simple test procedure to bypass the nasal swabs and the time-consuming PCR molecular amplification, currently being used by health professionals. Due to the long “silence” period of COVID-19 virus in our body, our health professionals and border quarantine measures are pressured to identify the asymptomatic but highly risky individuals. This challenges our ability in pushing the Limit of Detection to below ng/mL. This suggests needs in point of risk testings, for COVID-19 and many future ones.

**Figure 7.**
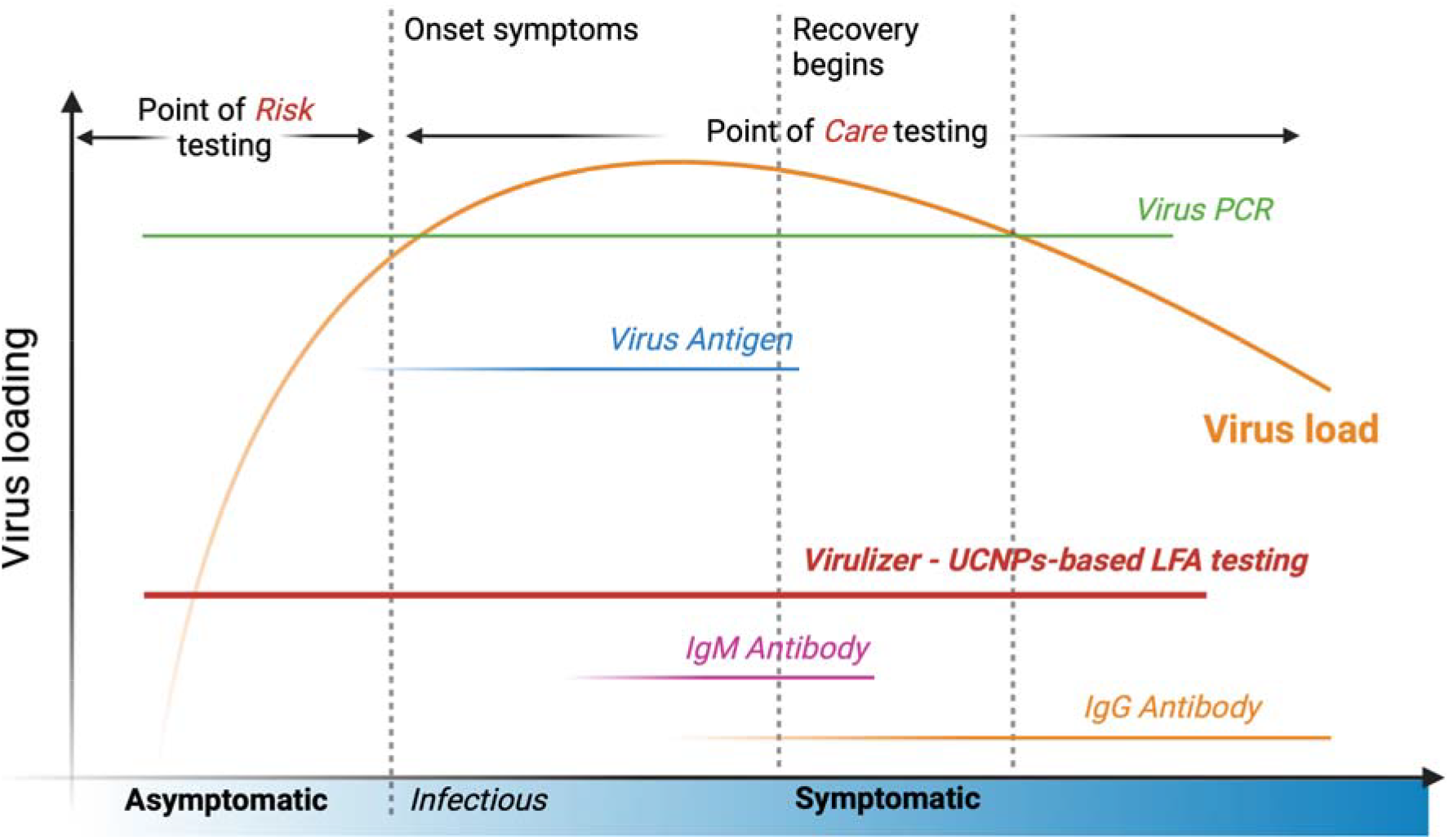
The methods and detection windows for COVID-19 along the virus load and clinical symptoms timeline. Current RAT and antibody testing (IgM/IgG) strips suffer from low sensitivity and are ineffective at detecting infections when viral loads are low, which limits early detection capabilities. The development of highly doped UCNPs-based LFA testing (Virulizer) significantly improves detection sensitivity to identify infections at the onset of infectiousness, rather than the onset of symptoms. Its ability to screen early-stage infections far surpasses that of traditional viral antigen and antibody tests, with sensitivity and accuracy comparable to PCR testing

RAT using UCNPs is promising in improving the LoDs and reduce the time to obtain results from several hours of PCR test to under 15 minutes. The saliva sampling and rapid testing methods together enhance testing efficiency, therefore to increase our capacity in the volume of tests per day, which is vital for managing current and future pandemics.

The broader application of UCNPs-based LFA tests extends beyond COVID-19, offering potential benefits in detecting and managing other infectious diseases, such as influenza and respiratory syncytial virus (RSV). Continuous research and development promise further improvements, driving the creation of advanced testing methods and procedures that contribute to better healthcare outcomes. The outlook includes the integration of AI-driven data analysis, which can predict outbreaks and guide public health responses more effectively. Moreover, this study underscores the importance of multidisciplinary and industry transformational collaborations among scientists, engineers, industry, healthcare professionals, and policymakers. Enhanced cooperation and knowledge sharing are essential for addressing a broad range of emerging infectious and health threats. Collaborative initiatives have already led to significant progress; for instance, international consortia have pooled resources and expertise to develop universal testing protocols and share real-time data on viral mutations.

In summary, this work represents a critical step forward in public health, with promising prospects for innovation and collaboration that will bolster our preparedness for future health crises. The combination of rapid, accurate testing, improved laboratory efficiency, and global cooperation forms a robust foundation for tackling future pandemics, ultimately leading to more resilient healthcare systems worldwide.

## Acknowledgements

We sincerely thank Professor William Rawlinson and Dr Gregory Walker from the University of New South Wales for their assistance with our work in the PC3 laboratory. This work was supported by the Australia China Science and Research Fund Joint Research Center for Point-of-Care Testing (ACSRF658277, SQ2017YFGH001190), Australian Research Council Industrial Research Hub for Integrated Device for End-User Analysis at Low-Levels (IH150100028), and the Australian Innovative Manufacturing Cooperative Research Centre fund (Rapid Point of Care SARS-CoV-2 Detection, Using A Sensitive Saliva Antigen Screening Test).

## Data and materials availability

All data are available in the main text and the supplementary materials.

